# Transformation of an olfactory placode-derived cell into one with stem cell characteristics by disrupting epigenetic barriers

**DOI:** 10.1101/2024.05.03.592460

**Authors:** Ghazia Abbas, Rutesh Vyas, Joyce C. Noble, Brian Lin, Robert P. Lane

**Affiliations:** Department of Molecular Biology and Biochemistry, Wesleyan University, Middletown, CT USA; Department of Developmental, Molecular and Chemical Biology, Tufts University School of Medicine, Boston, MA USA

**Keywords:** Epigenetics, Differentiation, Plasticity, Olfactory, Chromatin, Reprogramming

## Abstract

The mammalian olfactory neuronal lineage is regenerative, and accordingly, maintains a population of pluripotent cells that replenish olfactory sensory neurons and other olfactory cell types during the life of the animal. Moreover, in response to acute injury, the early transit amplifying cells along the olfactory sensory neuronal lineage are able to de-differentiate to shift resources in support of tissue restoration. In order to further explore plasticity of various cellular stages along the olfactory sensory neuronal lineage, we challenged the epigenetic stability of two olfactory placode-derived cell lines that model immature olfactory sensory neuronal stages. We found that perturbation of the *Ehmt2* chromatin modifier transformed the growth properties, morphology, and gene expression profiles towards states with several stem cell characteristics. This transformation was dependent on continued expression of the large T-antigen, and was enhanced by Sox2 over-expression. These findings may provide momentum for exploring inherent cellular plasticity within early cell types of the olfactory lineage, as well as potentially add to our knowledge of cellular reprogramming.

**SUMMARY STATEMENT:** Discovering how epigenetic modifications influence olfactory neuronal lineage plasticity offers insights into regenerative potential and cellular reprogramming.

## 1 INTRODUCTION

During early embryogenesis, pluripotent embryonic stem cells (ESCs) give rise to the various differentiated cell types of the adult body (Sanchez Alvarado and Yamanaka, 2014). This pluripotency is enabled by the expression of master regulatory transcription factors, such as Sox2, Nanog, Oct4, Klf4, and Myc (Chambers et al., 2003; Ellis et al., 2004; Loh et al., 2006; Masui et al., 2007), that preserve gene networks for mitotic activity and poised chromatin states, while suppressing the activity of master regulatory transcription factors required for differentiated cellular states (Oh and Jang, 2019). Differentiation of ESCs along a particular lineage and subsequent acquisition of a robust differentiated cellular state depends on genetic and epigenetic factors that entrench cellular networks appropriate for that cell type (Wutz, 2013). A dogma of cellular differentiation is the stability of cell states due to chromatin barriers that ensure irreversibility, as first proposed by Waddington’s landscape model (Waddington, 1957).

One of the important chromatin factors for establishment and maintenance of differentiated states is the Euchromatic Histone-lysine N-Methyltransferase 2 (EHMT2, also known as G9a, reviewed in (Shankar et al., 2013)) that is recruited to DNA through interactions with gene-specific targeting factors where it catalyzes the addition of local histone-3-lysine-9- methylation marks that lead to locally silenced chromatin (Tachibana et al., 2001). During early embryogenesis, G9a plays a critical role in cell differentiation by enabling developmental progression through silencing pluripotent factors (Chen et al., 2012; Lyons et al., 2014), as well as factors that enable alternative cellular states (Becker et al., 2016; Tachibana et al., 2002), and disruption of G9a has been shown to facilitate induced pluripotency (Rodriguez-Madoz et al., 2017). Thus, G9a is a central actor in the context of cellular differentiation, stability, and plasticity.

While there are a few striking examples of documented cellular dedifferentiation (Rajagopal and Stanger, 2016) that challenge the Waddington landscape model, these examples mostly represent atypical developmental contexts, such as tumor progression (Lee et al., 2016) or adult organ/tissue regeneration (Brawley and Matunis, 2004; Jopling et al., 2010; Kintner and Brockes, 1984; Shoshani and Zipori, 2011). One documented case of apparent dedifferentiation unfolds under a regenerative regime within the adult mammalian olfactory system. Given its vulnerability to environmental insult, various cell types of the olfactory epithelium (OE) must continuously undergo replenishment throughout adult life (Graziadei and Graziadei, 1979; Kondo et al., 2010; Mackay-Sim and Kittel, 1991), and as such, represents one of the few known examples of mammalian neural regeneration (Murray and Calof, 1999). During a regenerative regime, a population of mitotically active basal cells differentiates into both neuronal and/or non-neuronal cell types (Gadye et al., 2017), the former involving a progression to committed neural progenitors (dGBC cells), immature olfactory sensory neurons (iOSNs), and finally mature olfactory sensory neurons (mOSNs) (Graziadei and Graziadei, 1979). Strikingly, injury-induced dedifferentiation of committed neural progenitors into multipotent stem cells has been observed (Lin et al., 2017), underscoring an ability of OE cells to overcome epigenetic barriers in response to severe injury, presumably as a hedging strategy to allocate cellular resources as needed.

Our laboratory employs the olfactory placode- (OP-) cell lines, OP6 and OP27, both of which were clonally derived from developing E10.5 mouse embryos and immortalized using a temperature-sensitive Large-T antigen (Illing et al., 2002). These cell lines had previously been characterized as an intermediate stage of the olfactory sensory neuronal (OSN) lineage, and both lines can be induced by retinoic acid to further differentiate into neurons that express markers appropriate for mature OSNs (Illing et al., 2002; Pathak et al., 2009). In this study, we interfered with G9a protein function to investigate developmental consequences in these OP cell lines, inspired by previous observations made about G9a function during OSN progression (Fiszbein et al., 2016; Lyons et al., 2014; Roopra et al., 2004), as well as more generally in neural progenitor reprogramming (Ma et al., 2008). We found that G9a depletion caused rare transformation of OP cell types to adopt altered cell morphology and growth characteristics, as well as reorganized heterochromatin and up-regulation of genes associated with acquisition of pluripotency. These findings raise the possibility that differentiated cells along the olfactory neuronal lineage might retain atypical plasticity in support of regenerative processes, and thus, could represent an as yet untapped resource for neuronally-derived induced pluripotent stem cell (iPSC) protocols.

## 2 Results

We used two immortalized olfactory placode-derived cell lines, OP6 and OP27 (Illing et al., 2002), in order to investigate cell state stability in the developing olfactory system. These lines were previously characterized based on marker proteins whose expression profiles suggested a stage between a basal progenitor cell and mature olfactory sensory neuron: e.g., they are GAP43 positive (an immature OSN marker), yet OMP negative (a mature OSN marker) and Neurog1 negative (a committed neural progenitor cell marker) (Illing et al., 2002; Pathak et al., 2009). Both OP6 and OP27 cells can be induced with retinoic acid to differentiate into bipolar neurons that express more mature OSN marker proteins, such as G-Olf, OMP, FOXG1, O/E-1, and adenyl cyclase III, as well as exhibit bipolar extensions characteristic of mature OSNs in vivo (Pathak et al., 2009). Both lines also express olfactory receptor (OR) genes in a mutually exclusive manner (Abbas et al., 2021; Noble et al., 2018; Pathak et al., 2009) as is evident for immature and mature OSNs in vivo (Nagai et al., 2016; Serizawa et al., 2000). Based on these prior studies, we had inferred that these cell lines model an immature OSN (iOSN).

Despite these affirming phenotypic traits, the OP cell lines have not previously been characterized at the transcriptome level, where we anticipated that immortalization by the Large T-antigen would likely impose a large global impact on RNA expression profiles (Furuya et al., 2021). Here, we analyzed the bulk transcriptome of OP6 cell populations to gain further insights into the developmental status of this cell line. Indeed, we found that as compared to transcriptome profiles for native iOSN cell populations, numerous genes were >2-fold up- or down-regulated in OP6 relative to iOSNs (Fig. 1A). In consideration of just iOSN signature genes (Zunitch et al., 2023), OP6 cells tended to under-express at the RNA level (Fig. 1B). The log2 plot in Figure 1B showing signature-gene down-regulation in OP6 cells relative to iOSNs is strikingly similar to a previously published log2 plot comparing signature gene expression in primary versus immortalized endothelial cell cultures ((Deng et al., 2020); see Fig. 2B in their paper). Based on a previous study that reported genes that were significantly differentially expressed upon induction of the Large T-antigen (Furuya et al., 2021), we noted that more than half of the signature genes compiled for iOSNs, as well as other cell types from the olfactory epithelium (OE), were significantly impacted. The fact that immortalized OP6 cells nevertheless reliably exhibit phenotypic attributes of an iOSN suggests a robust system of post-transcriptional regulation to stabilize cell identity.

**Figure 1.**
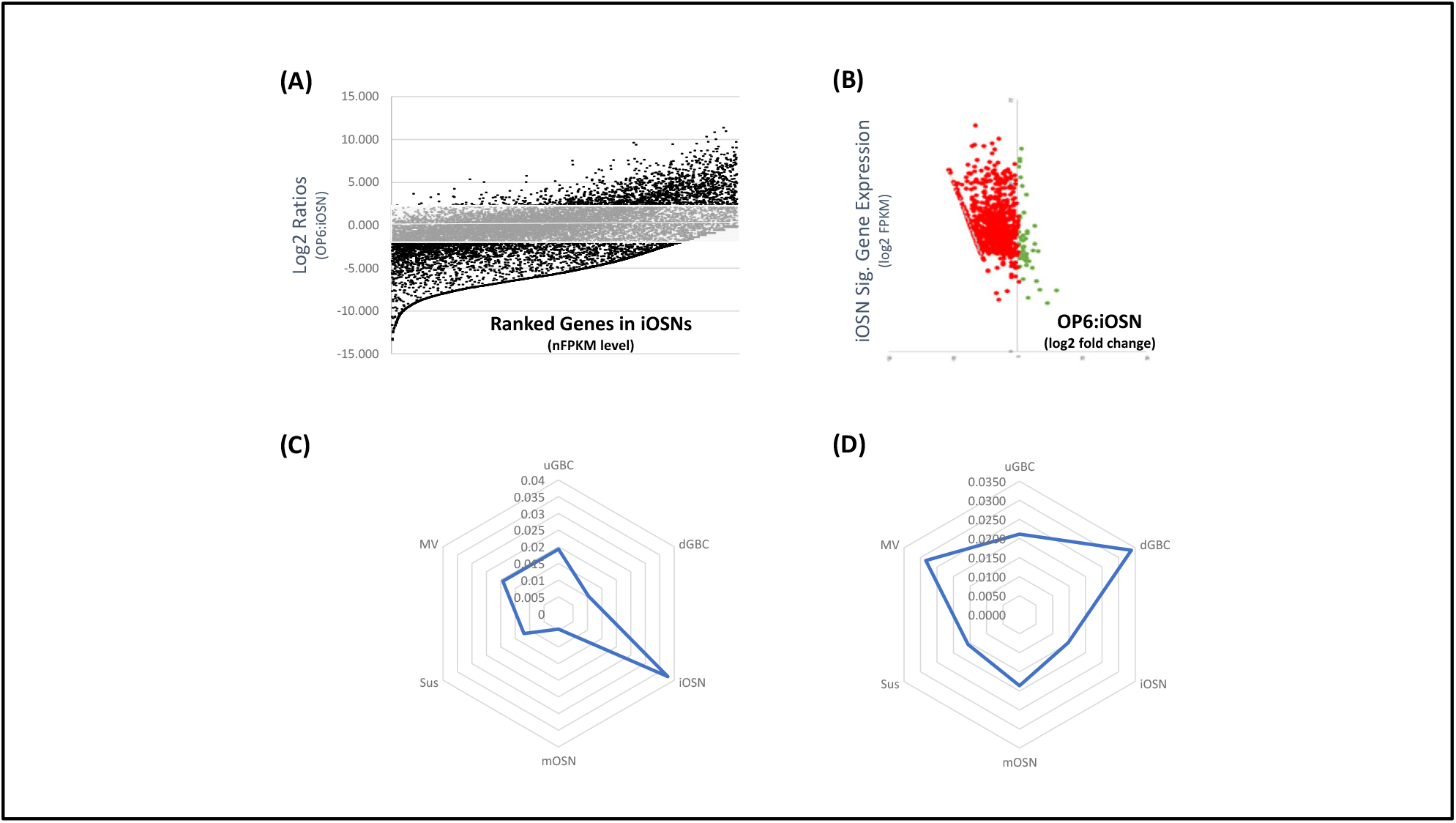
OP6 cells have diverged from iOSN RNA expression profiles. (A) Log2 ratios of nFPKM expression levels in bulk OP6 populations relative to ranked expression in pseudobulked iOSN cells. The rectangle captures all genes whose expression in OP6 is within twofold of iOSNs. **(B)** Plot of Log2 expression of 999 iOSN signature genes and corresponding Log2 change in OP6 cells relative to iOSN expression. **(C)** All misregulated genes with Large T- antigen expression were removed from each of the OE signature-gene lists, and these corrected lists were scored for average FPKM levels in OP6 cells weighted for relevance. (**D)** A set of 500 genes misregulated with Large T-antigen expression were scored as in C against each of the 6 ranked OE signature gene lists.

**Figure 2.**
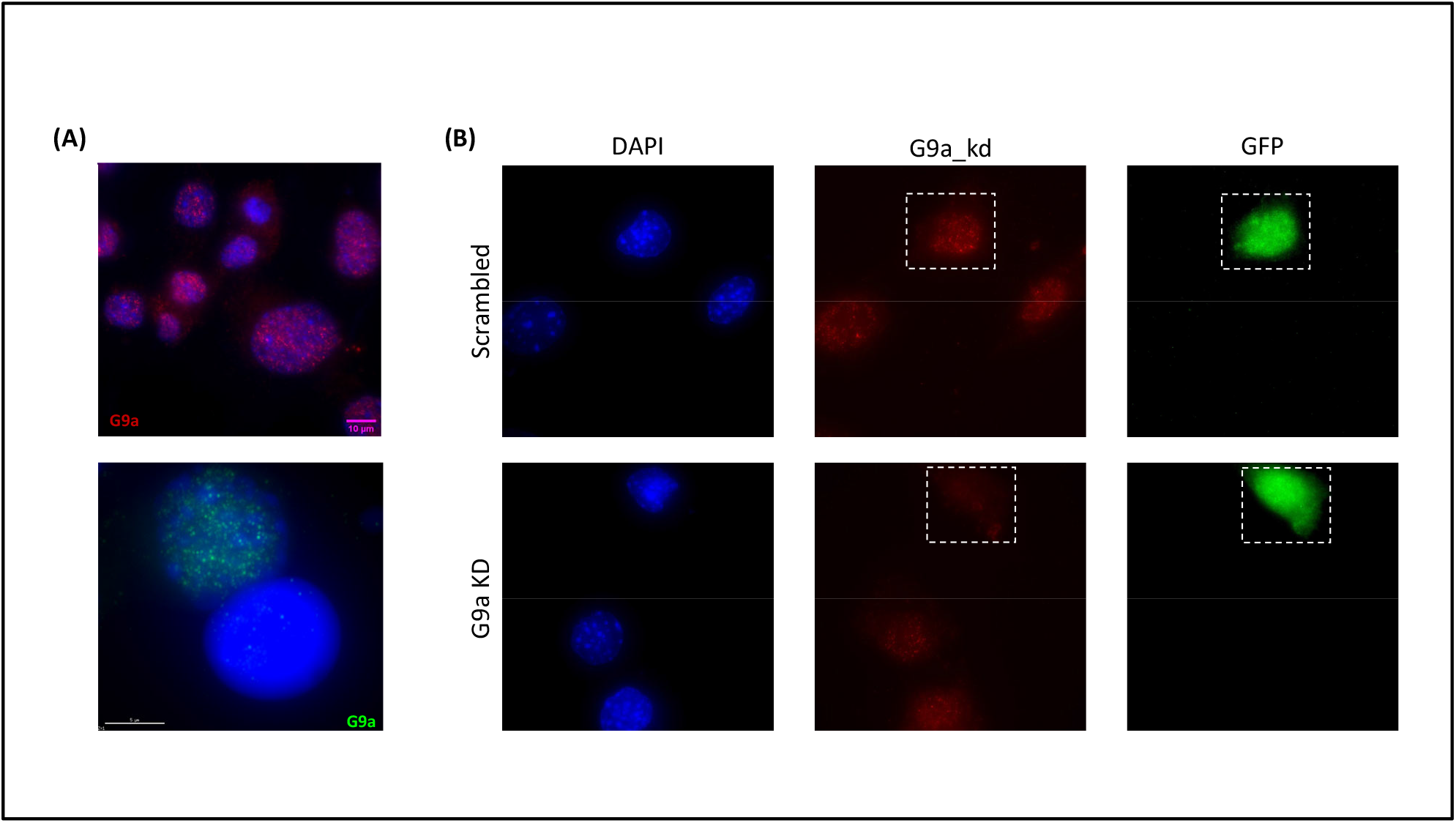
G9a Expression in Scrambled and GFP+ cells. (A) *Top:* G9a IF in untreated OP6 cells. *Bottom:* Example of two nearby cells following transfection with the G9a-targeted CRISPR-Cas9, with the bottom cell having knocked out G9a and the top cell being un- transfected OP6. Scale bar = 5 mm. **(B)** Representative images showing G9a expression in scrambled (top panel) and G9a knockout (bottom panel) samples. Hashed boxes highlight a GFP+ cell in both the scrambled and knockout populations. Quantitation of average G9a IF intensity in untreated OP6 populations versus CRISPR-treated populations is shown in Supplementary Figure 2.

We trimmed genes previously shown to be misregulated with Large T-antigen expression (Furuya et al., 2021) from the ranked set of signature genes in each of six different OE cell types (Zunitch et al., 2023). We scored OP6 cells based on the normalized FPKM expression level for the top 250 signature genes in each list weighted for relevance based on their ranked position (see Methods). A radar plot showing scores for each OE cell type indicated that OP6 cells exhibited more robust RNA expression of the most relevant iOSN signature genes as compared to other OE signature genes (Fig. 1C). Therefore, despite significant disruption due to immortalization, OP6 cells nevertheless exhibit transcriptome profiles consistent with an iOSN founder cell state. When using the same scoring system to assess only those Large T-antigen misregulated genes trimmed away in the above analysis, we found that these specific genes would tend to bias scoring towards a mitotically active dGBC cell type of the olfactory neuronal lineage (Fig. 1D).

### 2.1 Interference of H3K9 methylation in OP6 cells by knocking down G9a protein function

To investigate phenotypic stability of the OP lines, we interfered with a chromatin modifying factor, G9a, previously shown to be important in the development of olfactory sensory neurons. Using two separate perturbation protocols – either a CRISPR-Cas9 cassette or BIX01294 (Sigma) drug inhibitor – we perturbed G9a protein expression and function, respectively. The small-molecule inhibitor, BIX01294, is a drug that occupies the histone binding site of G9a in order to prevent interactions with histones and inhibit its methyl-transferase activity (Chang et al., 2009; Kubicek et al., 2007). As shown in Supplementary Fig. 1, this treatment significantly reduced H3K9 di- and tri-methylation levels in OP6 cells as measured by IF and Western blotting, consistent with a reduction in G9a protein function.

The CRISPR-Cas9 treatment targeted exon-8 near the beginning of the G9a gene. Following treatment, transcript analyses identified a cluster of disruptive base deletions that far exceeded mutation frequencies identified in scrambled controls (Suppl. Fig. 2A). Overall, ∼42% of RNA transcripts within G9a-targeted populations exhibited an insertion/deletion that would result in a disruptive frame-shift in the G9a gene. Such a result would be more consistent with the average targeted cell possessing a heterozygous as opposed to a homozygous deleterious genotype. Consistent with the interpretation that a full deleterious genotype was rare, we observed a distribution of non-zero levels of G9a protein expression levels in sorted G9a_kd cells; i.e., with none of the targeted cells sampled exhibiting completely abolished G9a protein expression relative to background (Suppl. Fig. 2B). The overall average G9a protein expression reduced to ∼48.5% of wildtype levels. Examples of G9a expression levels in knockdown versus scrambled control cells are shown in Figure 2.

We examined G9a transcript levels in the GFP-sorted sample in order to explore whether CRISPR treatment had resulted in a G9a RNA knockdown effect. Bulk sequencing exhibited a small average decrease in G9a RNA levels in two independent knockdown experiments (average=74.5% versus bulk scrambled controls), although this difference was not significant (paired T-test p=0.15). To gain further insight at higher resolution, we examined G9a transcript levels at the single-cell level. We found that the distribution of G9a transcript levels were heterogeneous and largely overlapping between the G9a_kd sample and control sample (Suppl. Fig, 2C). We speculate from these observations that the reduction in G9a protein expression in knockdown cells might have arisen from a combination of effects. For example, the subpopulation of GFP+ cells that exhibited normal G9a transcript levels presumably reduced G9a protein by non-productive translation (e.g., due to reading frame disruptions) or decreased protein stability (e.g., due to point mutations producing destabilizing amino-acid substitutions). On the other hand, GFP+ cells with significantly decreased G9a RNA levels might have produced a lower dosage of G9a protein due to CRISPR-mediated RNA knockdown effects, such as transcriptional steric interference (Larson et al., 2013), recruitment of heterochromatic factors to the targeted locus ((Anton and Bultmann, 2017; Lo and Qi, 2017) and/or nonsense-mediated RNA decay (Schoonenberg et al., 2018).

### 2.2 G9a knockdown resulted in a small global transcriptome shift

We calculated pairwise Pearson correlation coefficients (PCC) for principal components distinguishing sequenced samples, as well as scatterplot r^2^ values (RS) of overall gene expression levels, to assess whether scrambled control samples caused transcriptome shifts relative to untreated OP6 cells, as well as to assess reproducibility between two independent G9a-targeted CRISPR experiments. Both scrambled control replica experiments each exhibited RS values > 0.96 when compared with untreated OP6 cells, and exhibited PCC >+0.98 with each other (data not shown). Similarly, the two replica G9a-targeted experiments exhibited PCC > +0.98 with each other. These data indicated reliable reproducibility in both the CRISPR control and targeted experiments, and that the scrambled control was likely a good proxy for untreated OP6 cells. When comparing each G9a_kd sample to their corresponding scrambled control sample, the RS and PCC values were reduced by a similar magnitude in both experiments (the average PCC reduced from +0.985 to +0.974 in the two G9a_kd versus scrambled comparisons; T-test p-value=0.036). These observations indicated a small but measurable transcriptome impact with perturbation of the G9a gene. Overall, these two independent CRISPR-mediated G9a_kd experiments revealed a subset of ∼4% of differentially expressed genes (Suppl. Fig. 3A). Approximately the same number of genes were up-regulated as down-regulated, although the most significantly changed genes were more likely to be up-regulated in knockdown samples (Suppl. Fig. 3A).

### 2.3 Emergence of rare 3-dimensional colonies within G9a-perturbed OP cell populations

After several days of post-transfection growth in normal OP cell culture media, we observed rare formation of 3-dimensional colonies within G9a_kd transfected cultures (but not in scrambled controls) in both the OP6 and OP27 cultures (Fig. 3). The observed incidence was merely ∼3-5 colonies per million cells, or approximately ∼1 colony per 10,000 GFP+ cells (Fig. 3B). An example of colony growth over a 9-day period is shown in Figure 3A. Both OP6 and OP27 cell lines also exhibited rare colony formation with BIX01294 treatment (Fig. 3B). The morphological characteristics of the drug-induced colonies resembled colonies observed with the previous CRISPR treatment, suggesting that colony formation depended on the loss of G9a protein activity in both scenarios. In contrast, we never observed colony formation when treating the immortalized GD25 fibroblast cell line (Wennerberg et al., 1996) with either G9a- perturbation method (Fig. 3B), suggesting that this emergent property might have depended on factors present in cells derived from the olfactory lineage.

**Figure 3.**
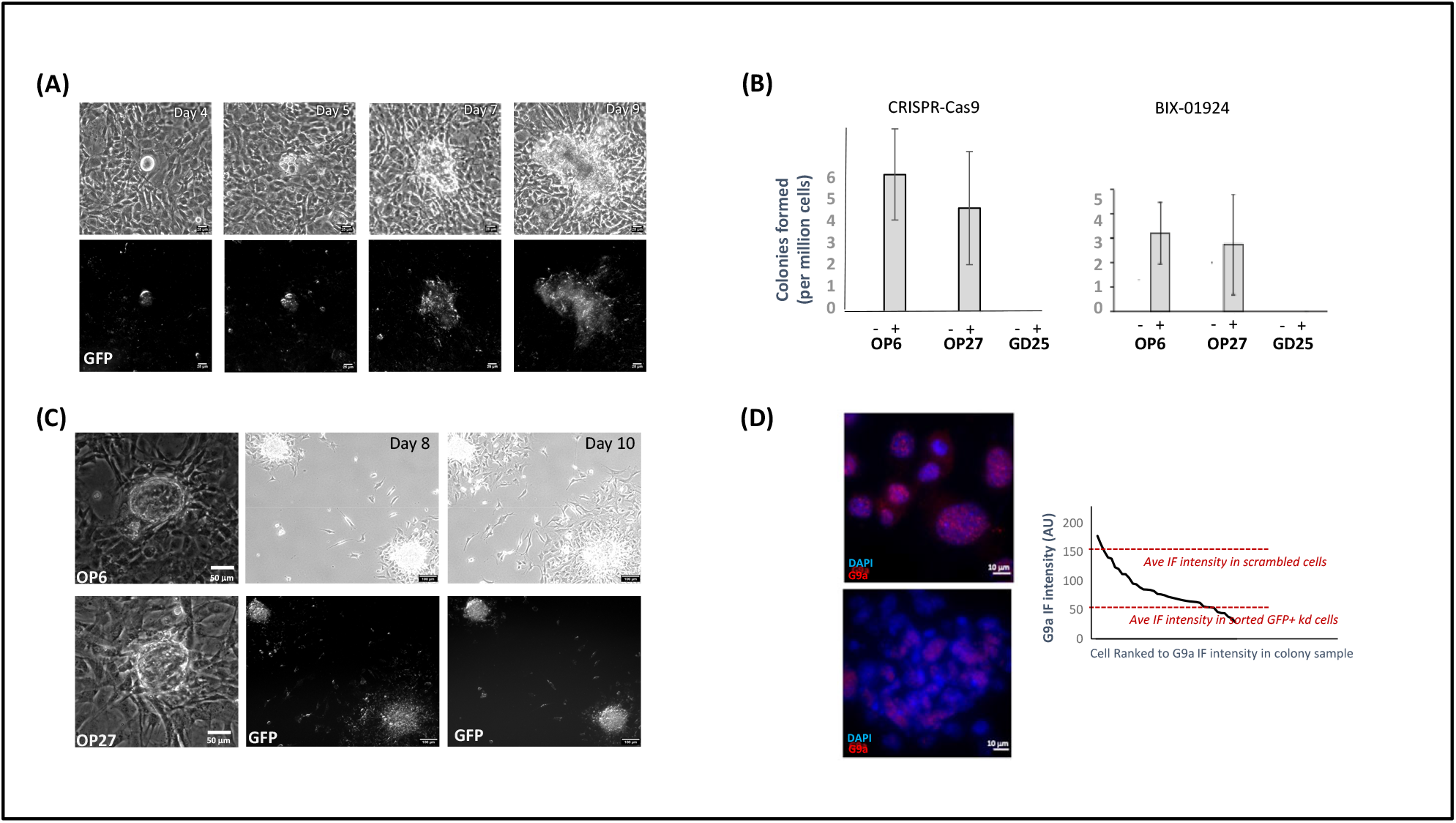
Rare colonies form in G9a_ko cultures that exhibit gene up-regulation. (A) Example of colony maturation from G9a_ko populations over a 9-day period. **(B)** Colony formation incidence in three different immortalized cell lines (OP6, OP27, GD25) with both CRISPR treatment and using the BIX-01924 G9a inhibitor drug. **(C)** *Left:* Example of a colony formed in treated OP6 (top) and OP27 (bottom) populations upon G9a inhibition with BIX- 01924. *Middle and Right:* A pair of secondary colonies produced from passaging a primary OP6 colony derived from CRISPR treatment at Days 8, and 10 post-passage, shown in both brightfield (top panels) and GFP fluorescence (bottom panels). **(D)** *Left:* Immunofluorescence (IF) staining of G9a (red) in untreated OP6 cells (top) and within colonies (bottom). Nuclei are counterstained with DAPI (blue). *Right:* The distribution of ranked G9a IF intensities within colony cells (n=40 random cells from four representative colonies). Average IF intensity within scrambled cells (upper line) and G9a_kd populations (lower line) are indicated.

OP6 colonies could be scraped from a primary CRISPR-transfected or BIX-treated plate and re-plated in secondary cultures, where these transplanted colonies continued to grow robustly (Fig. 3C). Interestingly however, when we broke up a colony into a monolayer of cells, we did not observe renewed colony assembly. This result hinted that initial colony formation might have required a cellular state present within the primary transfected culture, and that the individual colony cells were not autonomous colony-forming units. Moreover, when we sorted G9a-targeted GFP+ cells, we also did not observe colony formation even after prolonged culturing. This observation is consistent with the hypothesis that colony formation might have required a collaboration between wildtype and G9a knockdown cells, the former having been excluded from GFP+ sorted populations. Consistent with this hypothesis, we note that G9a expression, while reduced as compared to untreated OP6 populations, was surprisingly heterogeneous within colonies, including numerous cells with near or above wildtype levels of G9a protein expression (Fig. 3D). From these observations, we speculate that the presence of both G9a+ and G9a knockdown cells within colonies might reflect a colony-forming partnership between these two cellular states.

### 2.4 Colonies exhibit altered growth and cell morphological characteristics

OP6 cells typically grow in a monolayer on the bottom surface of culture dishes without the need of supporting cells or matrix; we have never before observed 3-dimensional cell communities form under our typical culturing conditions or arising from numerous previous genetic manipulations of this cell line. We made several phenotypic observations about these colonies. First, we observed that colony growth was accelerated relative to the growth of untreated OP6 cells, especially in younger/smaller colonies (Fig. 4A; left). Second, colony growth was sensitive to crowding, as evident by reduced and/or negative growth rates when a colony was grown at higher cell density (Fig. 4A, middle and right panel). Third, colonies could neither form in the first place, nor be maintained, when the Large T-antigen was deactivated at the non-permissive temperature (39°C), indicating that these colonies had not attained mitotic independence (Fig. 4B).

**Figure 4.**
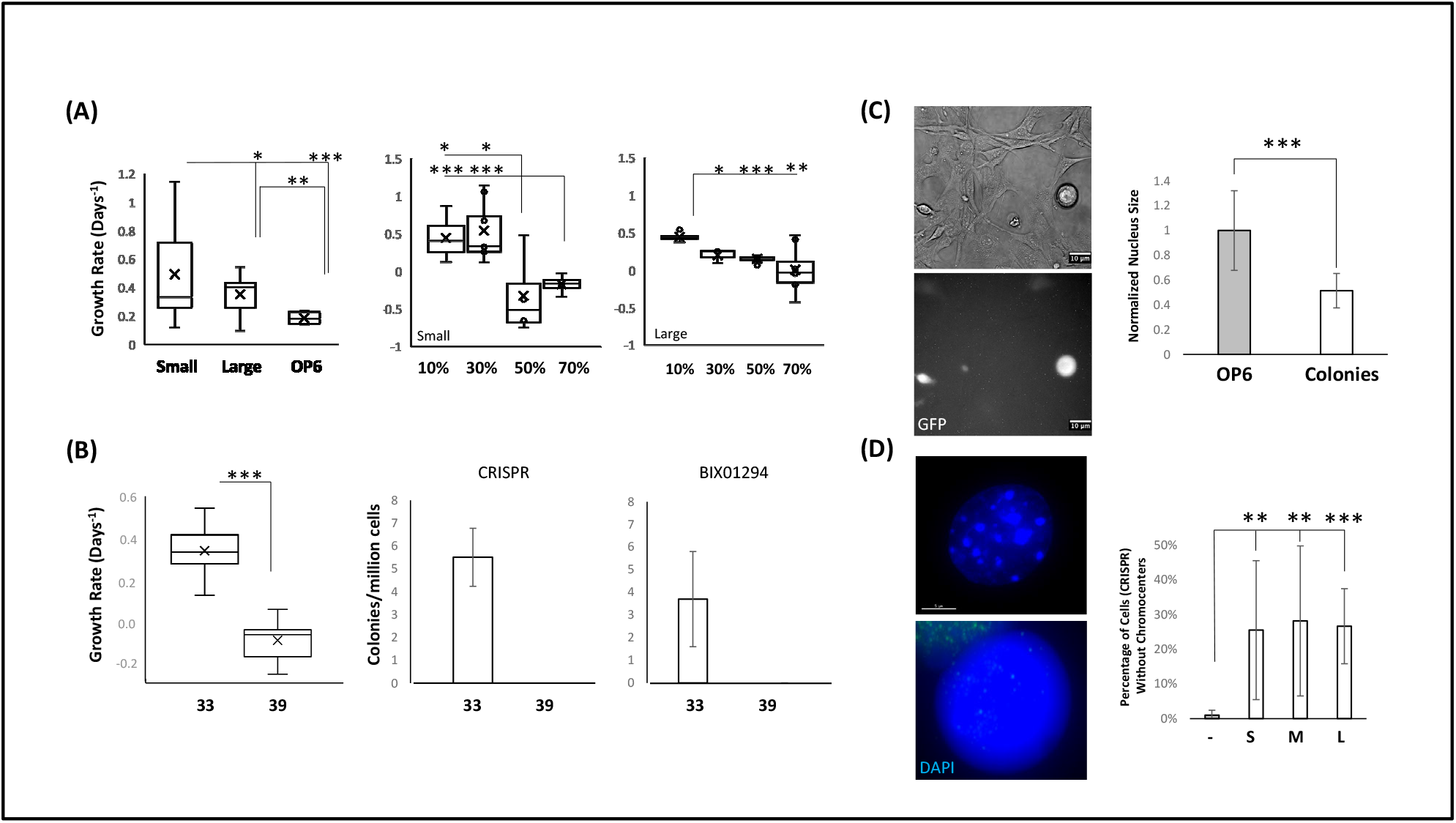
Colonies exhibit numerous altered cellular phenotypes. (A) *Left:* Growth rates are increased in small and large colonies relative to untreated OP6 cells. *Middle and Right:* Growth rates for small and large colonies at varying surrounding cell densities. **(B)** Colony growth rates (left) and colony formation (middle and right) cease when the Large T-antigen is deactivated at the non-permissive temperature (39^0^ C). **(C)** *Left:* GFP+ cells commonly exhibit a more rounded cell shape relative to background non-transfected/GFP- cells. *Right:* G9a-perturbed OP6 colonies exhibit reduced nucleus size. **(D)** *Left:* The distribution of DAPI-dense staining becomes more uniform with apparent depletion of nuclear chromocenters. *Right:* Quantitation of the percentage of cells in untreated (-), small (S), medium (M), and large (L) colonies that lack chromocenters.

The physical transformation of OP6 cells apparent within colonies was quite dramatic in several additional ways. Fourth, although there was heterogeneity, GFP-positive cells tended to be more compacted and spherical in shape (Fig. 4C, left panels). Compacted, rounded cell shapes have been associated with a less differentiated state, including undifferentiated embryonic stem cells (ESCs) (Bongiorno et al., 2018). Fifth, we observed a reduction in average nucleus size (Fig. 4C, right panel). Reduced nucleus size has been associated with undifferentiated ESCs, certain types of tumor cells, as well as micronuclei arising from kinetochore or other genotoxic defects (Butler et al., 2009; Koike et al., 2007; Ye et al., 2019). Sixth, colony cells tended to exhibit a change in heterochromatin organization, with a reduction of DAPI-dense chromocenters (Fig. 4D); since chromocenter formation is dependent on H3K9me3 levels (Harničarová Horáková et al., 2010), this observation was consistent with a loss of G9a-mediated maintenance of these marks. We note that diffuse and disorganized heterochromatin has been associated with pluripotent stem cells (de Wit et al., 2013; Fussner et al., 2011; Meshorer et al., 2006; Wijchers et al., 2015). Overall, these transformations in cell morphology and growth properties provided initial momentum to explore whether G9a perturbation had caused OP6 regression to an earlier developmental stage.

### 2.5 Colonies up-regulated genes associated with cell communication, cell motility, cell proliferation, and various developmental processes

Bulk sequencing revealed a large set of up- and down-regulated genes in both small and large colonies (Suppl. Fig. 3B). A preliminary Gene Ontology (GO) analysis identified functional enrichments in sorted G9a_kd cells and colonies versus scrambled control cells that would seem supportive of *de novo* colony organization (Suppl. Fig. 4A). Notably, enrichment categories included cellular movement and migration, as evidenced by terms such as “positive regulation of cell motility,” and “chemotaxis.” Moreover, there was enrichment in terms related to vascular formation, angiogenesis, tissue remodeling, and organogenesis. Additionally, genes implicated in extracellular matrix (ECM) biology were enriched, encompassing terms like “extracellular matrix component,” “collagen binding,” and “integrin binding,” underscoring potential roles in colony cellular adhesion, signaling, and tissue structure maintenance. Furthermore, receptor-related terms signified potential enhancement of signal transduction and cellular communication processes. Together, these GO terms associated with colony formation suggests a complex and diverse molecular milieu, with the up-regulation of specific processes that might be anticipated with changes in cell-cell interactions and developmental status.

Since we had determined that colonies were heterogeneous based on G9a expression status (Fig. 3), we were next interested in looking at transcriptome heterogeneity within these “transformed” cell populations, comparing colony cells to ancestral G9a_kd cells, as well as scrambled control cells, at a single-cell resolution. Because of the stringent extracellular matrix within colonies where single cell isolation would have required very harsh treatments, we deemed it impossible to utilize a typical cytoplasmic-based single-cell RNA sequencing pipeline, and instead utilized a nucleus-based RNA isolation strategy commonly employed for hard tissue samples (Guo et al., 2023). Since we were aware that nucleus-derived RNA differs in substantial ways from cytoplasmic-derived RNA (Ding et al., 2020), we conducted the same nucleus isolation protocol on sorted GFP+ G9a_kd cells, as well as from a mixture of scrambled and untreated OP6 controls. Our initial analysis was based on the routine bioinformatics package purchased through Novogene, the company that generated the sequencing data for us. As shown in Supplementary Fig. 4, the integrated single-cell dataset included two clusters (Novogene clusters 3 and 8) that had ∼homogeneous colony cell compositions (Suppl. Fig. 4C, 4D). Among the most significantly up-regulated genes in these two colony-dominated clusters were Cyclin Dependent Kinase 8 (CDK8), Calcium/Calmodulin Dependent Protein Kinase ID (Camk1d), and translation-associated genes Cmss1 and Lars2, as well as the Metallothionein 1 (Mt1) gene and the GM42418 cell differentiation lincRNA. The GM42418 gene is a marker for hepatocyte cell differentiation (Han et al., 2022) and the Mt1 gene encodes a metal-binding protein that has been closely linked with the regulation of cell proliferation and immune cell differentiation (Dai et al., 2021). The other four genes (CDK8, Camk1d, Cmss1, and Lars2) are putative factors that regulate pluripotent embryonic stem cells (ESCs) (Adler et al., 2012; Bergamaschi et al., 2008; Feng et al., 2022; Wang and Amoyel, 2022). We will elaborate on these latter inferences in the context of discussing other ESC-like features of OP6-derived colonies in a subsequent section.

### 2.6 Analysis of sample heterogeneity and transitions between subpopulations

We conducted our own bioinformatics on the single-cell RNA-seq dataset in order to ask a set of specific questions relating to our study. We initially clustered without integration in order to more cleanly assess cell heterogeneity within particular samples (Fig. 5A). Within the control sample (scrambled and untreated OP6), we observed 4 clusters (C0=∼54% of sample, C2=∼30% of sample, C6=∼8% of sample, and C7=∼7% of sample). Some of the observed heterogeneity could be predicted by cell cycle stage. For example, 99% of C2 cells were going through the cell cycle (85% at the G2/M stage; 14% in S-phase), and to lesser extent, 93% of C7 cells were likewise in S-phase or at G2/M, whereas the C0 and C6 clusters had a much higher percentage of cells in G1. (Fig. 5B). We also presume that some of the heterogeneity evident within control OP6 populations might be partly explained by mutational histories – e.g., due to aneuploidies that can commonly arise with prolonged culturing of a cell line (Devalle et al., 2012). In addition, some of the observed heterogeneity is likely due to the presence of both scrambled CRISPR+ cells and untreated CRISPR- cells in this control sample, where retroviral infection even with a scrambled/un-targeted guide RNA would likely have impacted the transcriptome (Moiani et al., 2012). We note that preliminary RNA velocity analyses suggested that transitions between subpopulations dominated by cells in S- or G2/M-phase of the cell cycle (C2 and C7) to the other subpopulations with larger percentages of cells in G1 (Suppl. Fig. 5). Such transitions between subpopulations would seem to be more consistent with a common community of cells undergoing state changes at various point in time, as opposed to distinct cell communities stably differentiated by different mutational backgrounds.

**Figure 5.**
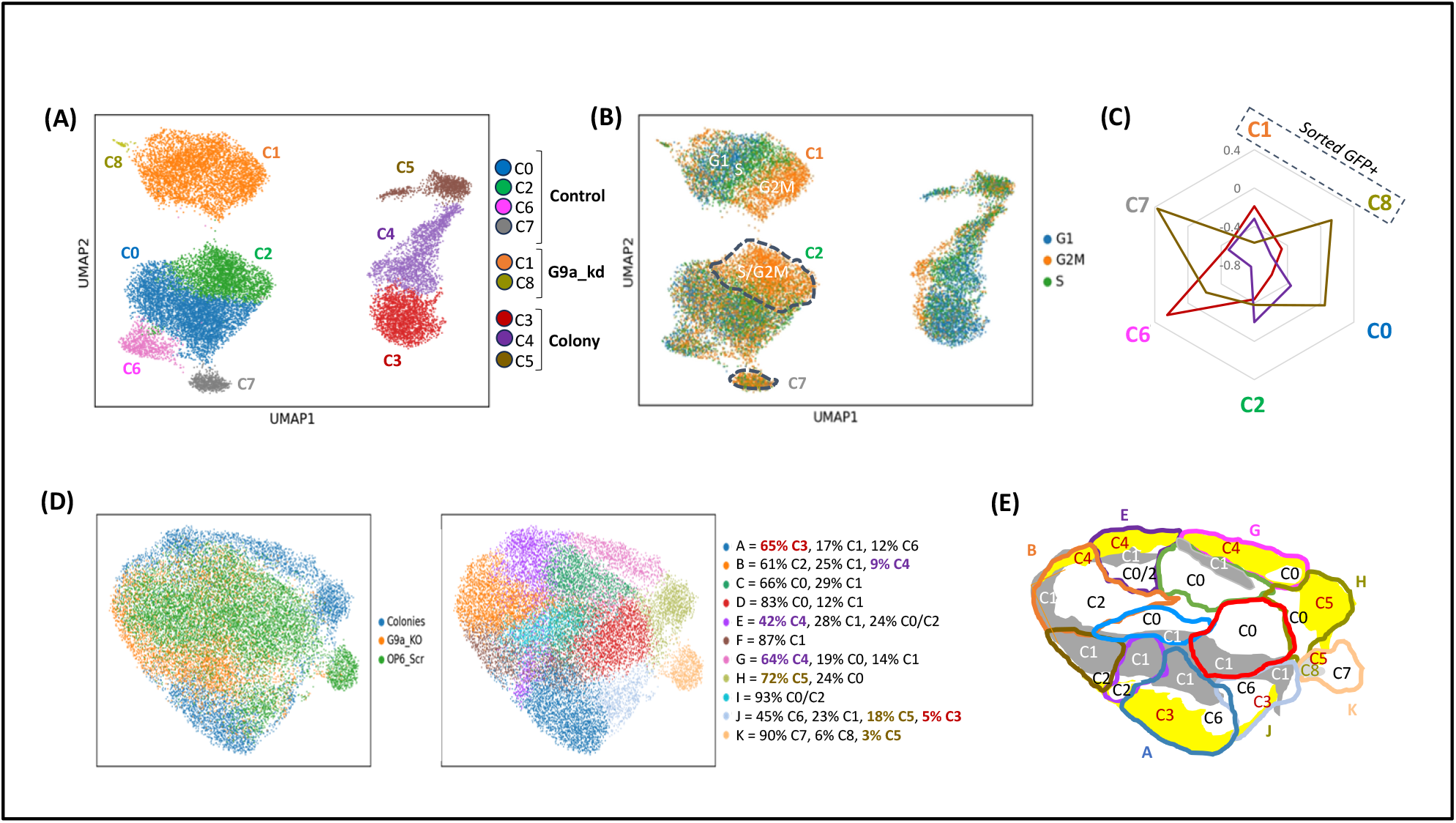
Single-cell analysis defines three distinct colony cell clusters each with correlated gene expression one of the ancestral subpopulations. (A) *Left:* Non-integrated UMAP plots showing control subpopulations (C0,C2,C6,C7), sorted G9a_kd subpopulations (C1,C8), and colony cell subpopulations (C3,C4,C5). **(B)** Non-integrated clusters are labeled for cell-cycle stage, including C2 (99%) and C7 (93%) control clusters dominated by cells going through the cell cycle. **(C)** Radar plots of PCC values comparing colony clusters (C3, C4, C5) to control (C0, C2, C6, C7) and sorted GFP+ (C1,C8) clusters. **(D)** UMAP plot of integrated single-cell data produces 11 clusters (A-K); non-integrated cluster composition is summarized in the table. **(E)** Representation of the integrated cluster map colored to indicate control cells (white), G9a_kd cells (gray), and colony cells (yellow) labeled with cluster numbers (C0-C8) corresponding to the non-integrated UMAP in panel (A).

The two G9a_kd cell clusters evident by single-cell RNA sequencing datasets included a majority C1 subpopulation (∼99% of sample) and a minority C8 subpopulation (∼1% of sample) (Fig. 5A). Stratification of the majority C1 cluster also appeared to correlate with cell-cycle stage (Fig. 5B). Given its tiny size, we wondered whether the C8 subpopulation might have been untransfected contaminants during the sorting process. To assess this, we identified a set of 74 significantly up-regulated genes and 31 significantly down-regulated genes that were >10-fold higher and lower, respectively, within bulk G9a_kd versus scrambled control samples (from two independent bulk sequencing datasets), and with a minimal FPKM=1.0 in the higher expressing sample. We found that both G9a_kd clusters (C1 and C8) exhibited a similar pattern of downstream gene up- and down-regulation (Suppl. Fig. 6A). Interestingly, RNA velocity analyses suggested transitions occurring from the C8 subpopulation to the C1 state (Suppl. Fig. 5), and thus, the tiny size of the C8 subpopulation might have been the consequence of transitioning to a more stable C1-like state following G9a perturbation.

The colony sample comprised three distinct clusters: C3=41% of sample, C4=38% of sample, and C5=21% of sample (Fig. 5A). We analyzed these 3 colony subpopulations to assess whether they each exhibited expected signatures associated with colony formation as had been evident from bulk sequencing datasets. We identified a set of 104 genes that had exhibited significant up-regulation in bulk colony samples versus G9a_kd samples, as well as a set of 229 genes up-regulated in association with bulk colony GO categories. In both cases, we found that all three colony cluster subpopulations exhibited higher average expression of these gene sets relative to other non-colony clusters, consistent with the interpretation that all 3 single-cell colony clusters had undergone “colony-like” transitions divergent from the wildtype state (Suppl. Fig. 6B, 6C). Interestingly however, we note that all 3 colony clusters were not “G9a_kd-like” in consideration of the up- and down-regulated genes identified in bulk G9a_kd vs control experiments (Suppl. Fig. 6A). This result was surprising because we knew that a large percentage of cells within colonies were G9a knockdown cells based on IF experiments (Fig. 3), and thus this observation raises the question as to whether there existed selective pressure for some degree of rescue of the downstream G9a_kd effects in order to favor colony growth or survival.

Finally, given that colonies comprised a mixture of cells with wildtype-like and G9a_kd expression as evident by IF experiments (Fig. 3), we assessed the 3 colony subpopulations for their transcriptome similarities with wildtype and G9a-targeted samples. We found that colony C5 cells were most well correlated with control OP6 cells and the least well correlated with the G9a_kd cells, whereas colony C3 cells were most well correlated with G9a_kd cells (Suppl. Fig. 6D). Colony C4 cells were maximally diverged from the original OP6 population (Suppl. Fig. 6D). We also note that this C5 subpopulation (most wildtype-like) appeared to be transitioning to the C4 colony state (least wildtype-like), as evident by RNA velocity analyses (Suppl. Fig. 5).

### 2.7 Single-cell sequence analyses indicate non-redundant relationships between each colony cluster and a corresponding ancestral subpopulation

We further analyzed Pearson Correlation Coefficients (PCC) and scatterplot r^2^ (RS) values to ask whether there existed relationships among the colony clusters, as well as with ancestral subpopulations (Suppl. Fig. 7A, 7B). The radar plot in Figure 5C is useful in visualizing our main conclusion. The colony C3 cells were best correlated with control C6 and the majority C1 knockdown clusters, whereas colony C5 cells were best correlated with the control C7 and the minority C8 knockdown clusters. The colony C4 cells were more ambiguous with respect to ancestral relationships. According to PCC values, C4 was only positively correlated with the other two colony subpopulations (C5 in particular), and exhibited no apparent relationship with any of the control subpopulations. On the other hand, according to RS values, C4 showed a close relationship with the C0/C2 control clusters, as well as the C1 knockdown cluster (Suppl. Fig. 7B).

When we mapped the non-integrated cluster identities as described in Figure 5A onto the integrated clustering evident in Figure 5D, we observed co-clustering of control and colony cells that mirrored inferences made from PCC and RS correlations discussed above (Fig. 5E): colony C3 cells co-clustered with control C6 cells, along with G9a_kd C1 cells, within the lower bulb of the integrated map (integrated clusters A&J); colony C4 cells co-clustered with control C0 cells, along with G9a_kd C1 cells, within the upper rim of the integrated map (integrated clusters BE&G), and colony C5 cells co-clustered with control C7 cells and G9a_kd C8 cells, within the lower right bulb of the integrated map (integrated cluster K). The only exception was integrated cluster H, which showed co-clustering of the largest fraction of colony C5 cells along with control C0 cells, without co-clustering with any of the G9a knockdown cells.

In sum, both the integrated and non-integrated data analyses suggested one-to-one relationships between specific subpopulations of colony cells and distinct ancestral subpopulations. We incorporate these apparent relationships into a model for colony formation from ancestral subpopulations within the Discussion section below.

### 2.8 Colonies subpopulations may have dedifferentiated to embryonic stem cell (ESC)-like states

We made three additional observations about transformed OP6 colonies that are noteworthy because they are also features of embryonic stem cell (ESC) colonies (Koike et al., 2007). First, transformed colonies, whether they are derived from CRISPR-mediated G9a knockdown or from BIX-mediated inhibition of G9a protein function, showed elevated staining of all five of the tested canonical marker proteins used to define pluripotent stem cell states (Fig. 6). Second, these colonies stained strongly for alkaline phosphatase (Fig. 6D), a marker for mouse ESCs (Andrews et al., 1984; Thomson et al., 1998). Third, more mature (larger) colonies transplanted and grown separately in secondary cultures often exhibited heterogeneity in cell morphology, with more compacted, rounder (and GFP+) cells evident at the core of the 3D mass, but also exhibiting a halo of more flattened cell monolayers at the periphery (Fig. 3C) reminiscent of embryoid bodies where differentiation occurs at the peripheral edges of stem-cell derived colonies (Rosowski et al., 2015). We also noted the down-regulation of the Pax6 neural lineage marker gene within colonies relative to ancestral G9a_kd cells and scrambled controls (Fig. 6B, 6C), which could signal dedifferentiation from pre-existing neuronal states.

**Figure 6.**
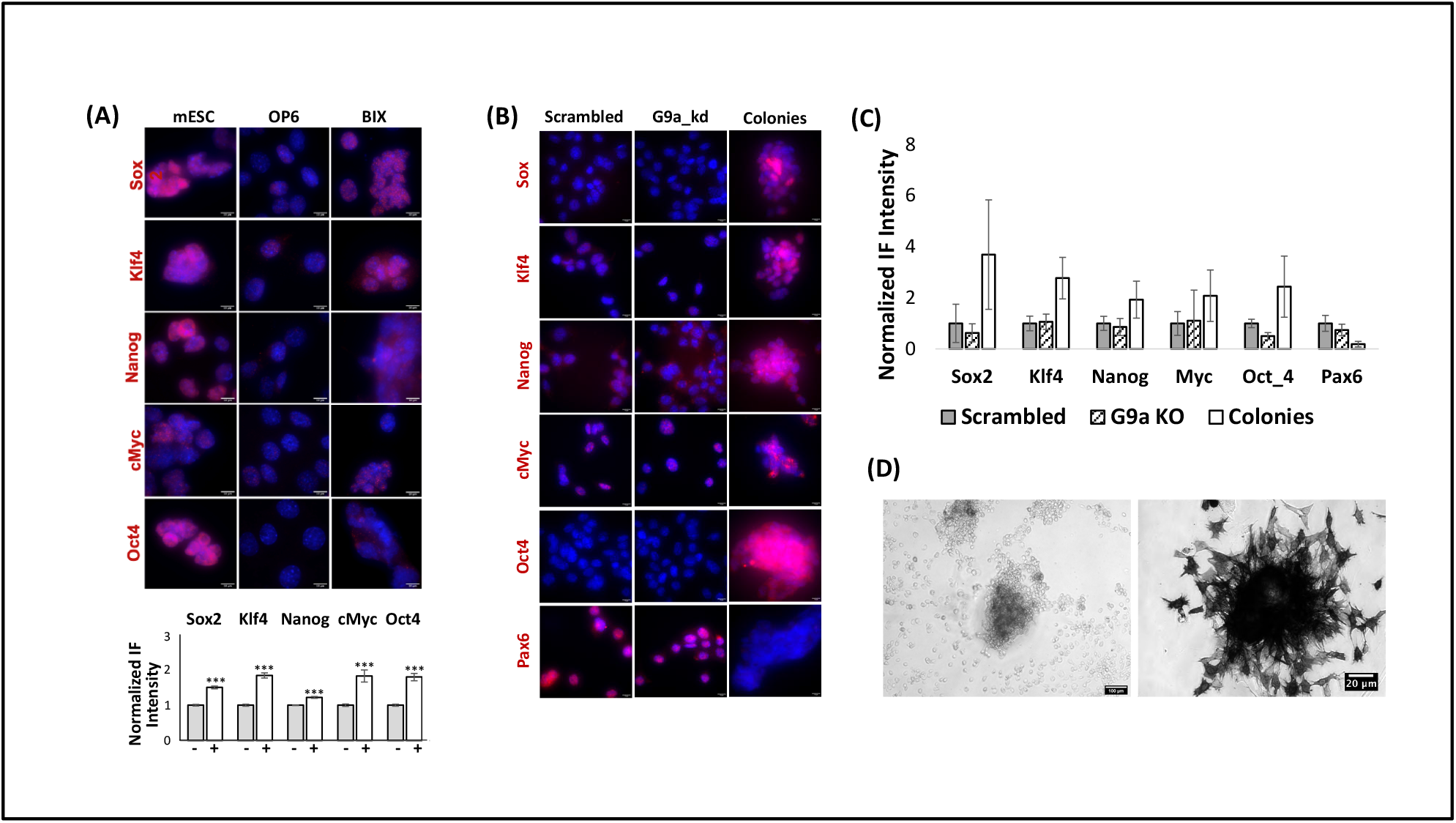
Immunofluorescence levels of canonical stem cell markers in G9a-perturbed OP6 colonies. (A) Immunofluorescence images of mouse embryonic stem cells (mESC) untreated OP6 cells, and BIX-treated OP6 colonies showing the expression of Sox2, Klf4, Nanog, cMyc, and Oct4 stem cell markers. Cell nuclei are visualized with DAPI staining (blue). Scale bars = 10 μm. *Lower*: Bar graph depicting the fold-increases of Sox2, Klf4, Nanog, cMyc, and Oct4 within BIX-treated OP6 cells (+) relative to untreated negative controls (-). **(B)** Up-regulation of stem cell markers Sox2, Klf4, Nanog, cMyc, and Oct4 in CRISPR-derived OP6 colonies compared to Scrambled Control and untransformed GFP+ cells. Cell nuclei are visualized with DAPI staining (blue). Scale bars = 10 μm. **(C)** Bar graph summarizing the normalized immunofluorescence (IF) intensities of stem cell markers from the analysis of several CRISPR-derived OP6 colonies (+) compared to Scrambled Control and untransformed GFP+ cells, as shown in (A). **(D)** Colonies stain with alkaline phosphatase.

As noted previously, the combination of using an immortalized cell line (with broad gene misregulation caused by Large T-antigen expression) and isolating RNA from the nucleus (with broad enrichment biases in comparison to cytoplasmic isolations) precluded confident mapping of colony transcriptome profiles to previously published ESC transcriptome profiles. Nevertheless, we attempted to find signal embedded within this anticipated noise to ask whether any of the three subpopulations of colony cells (C3, C4, or C5) seem to have transitioned towards a more ESC-like state at the transcriptome level. We first calculated r^2^ values for pseudobulked single-cell transcriptomes of each of the colony clusters as compared to a published bulk transcriptome of mouse ESC cells (Kinoshita et al., 2021). The strongest correlations were evident between the colony C4 subpopulation (Suppl. Fig. 8A), as well as the three C4-containing integrated clusters (G, B, and E; Suppl. Fig. 8B), whereas the colony C3 and C5 subpopulations were poorly correlated. We note that both the C2 (control) and C1 (G9a_kd) subpopulations also exhibited positive correlation with ESC transcriptomes (Suppl. Fig, 8A), the former presumably due to cell cycle gene up-regulation (99% of C2 cells are cell cycling) and the latter possibly due to up-regulation of genes that prime future colony transitions.

We next assessed apparent transitions towards ESC-like states versus away from iOSN- like states within various colony subpopulations. We first identified sets of genes exhibiting >10- fold up-regulation in published ESC (Kinoshita et al., 2021) versus iOSN datasets (Zunitch et al., 2023), and vice versa, genes exhibiting >10-fold up-regulation in iOSN versus ESC populations. We then calculated the average FPKM levels for these genes in each of the colony and non-colony cell clusters, as well as a net FPKM trajectory score reflecting the difference between ESC versus iOSN signatures. Based on this type of analysis, the colony C4 cluster exhibited the maximum positive ESC trajectory and the minimum iOSN trajectory (Suppl. Fig. 8C), consistent with the hypothesis that among colony cell populations, C4 cells appeared to have most transitioned away from iOSN profiles and towards ESC profiles. The same tentative conclusion was supported by analysis of iOSN signature genes, which were expressed at lower average levels in C4 cells as compared to other colony cells (Suppl. Fig. 8D). In contrast, the colony C5 cluster was least ESC-like (Suppl. Fig. 8A) and most iOSN-like (Suppl. Fig. 8C, 8D) based on these scoring methods, perhaps reinforcing the hypothesis that C5 cells were the most likely of the three colony clusters to have derived from wildtype ancestors.

In addition, we examined expression of a set of key marker genes within each of the colony clusters to further assess ESC-like characteristics. Transcript levels for three of the five canonical ESC marker genes, Sox2, Oct4 and Nanog, were below threshold in all three of the scRNA-seq clusters (C3, C4, and C5); since we know that the three encoded proteins for these genes were robustly expressed in colonies, the absence of above-threshold transcript underscores the fact that transcriptome analyses can be misleading with respect to steady-state protein levels and biological function within cells, especially with regard to low-expressing transcripts or transcription factors vulnerable to drop-out during single-cell RNA sequencing (L. Lun et al., 2016; Vallejos et al., 2017). Transcripts for the other two canonical ESC marker genes, Klf4 and c-Myc, were detected, and these two markers were expressed at maximum levels within colony C4 cells (Suppl. Fig. 8E). Colony C4 cells also expressed the highest levels of CDK8 and CAMK1D (Suppl Fig. 8E), the former involved with stem cell maintenance and self-renewal capacity (Adler et al., 2012; Fukasawa et al., 2021), and the latter involved in cell proliferation and epithelial-mesenchymal transitions (Bergamaschi et al., 2008). In consideration of our entire analysis, C4 cells would seem to be the best candidates within colonies to be responsible for the up-regulation of ESC marker proteins shown in Figure 6.

Finally, we note that a key signaling molecule used within stem cell niches, TGFB1 (Xu et al., 2018), was up-regulated in bulk colony cells versus controls, and was maximally expressed in colony C3 cells. Of the three colony clusters, C3 cells exhibited the strongest PCC values with G9a knockdown cells, and like C4 colony cells, were significantly diverged from wildtype OP6 cells (Suppl. Fig. 6D). We speculate on a colony-forming model in which the knockdown of G9a within these C3 colony cells might have resulted in the up-regulation of TGFB1, which in turn, signaled nearby wildtype-like C5 colony cells to diverge towards a more ESC-like state represented by colony C4 cells. We will consider this colony maturation model in the context of all of our data within the Discussion section below.

### 2.9 Sox2 over-expression synergistically contributes to OP6 colony formation

The low incidence of colony formation among a population of G9a-perturbed cells suggests a stochastic attribute that enabled transformation only under rare circumstances. One plausible stochastic attribute already discussed is the presumed rare occurrence of two nearby cells that differed in their G9a expression status forming a partnership in colony development. Another possible stochastic variable within populations could be the dynamic range of G9a perturbation due to variable penetrance of CRISPR-mediated mutations or local differences in BIX01294 drug efficacy. These differences may couple with other intrinsic states that could also be stochastic in nature, such as position in the cell cycle or the length of time spent during a critical stage of the cell cycle, factors known to be influential in reprogramming efficacy (Smith et al., 2010).

We speculated on another potential source of stochastic variability that might have been important for enabling colony transformation only in rare circumstances. As seen in Figure 7, most untreated OP6 cells expressed low levels of the Sox2 protein (if at all). Sox2 is a critical factor in OSN development (Gadye et al., 2017; Guo et al., 2010; Tucker et al., 2010), as well as in establishing and maintaining pluripotency of stem cell populations (Ellis et al., 2004; Masui et al., 2007). Sox2 expression evident in G9a_kd cells also exhibited a very low average level. However, we note that both untreated OP6 and G9a_kd populations exhibit rare outliers with elevated Sox2 expression that approximate the higher average Sox2 expression within colonies (Fig. 7), raising the question as to whether these outliers represented a rare prerequisite state for colony formation. To investigate this question, we induced two levels of Sox2 over-expression within OP6 cells. We found that colonies formed with Sox2 over-expression alone; i.e., even in populations that were not treated with G9a CRISPR-Cas9 knockdown (Fig. 7B). When combining Sox2 over-expression with CRISPR-mediated G9a perturbation, colony formation was significantly enhanced relative to either treatment alone. For example, within the “high” Sox2 overexpressing populations without G9a knockdown, as well as within G9a_kd populations without Sox2 over-expression, the incidence of colony formation was approximately 10 colonies per million cells in each case, whereas the combination of both treatments produced approximately 35 (3.5-fold more) colonies per million cells (Fig. 7B). These data, along with data already discussed, suggest that OP6 colony formation was enhanced by three conditions: expression of the Large T-antigen (i.e., mitotically active), elevated expression of the Sox2 transcription factor, and reduction of chromatin barriers through G9a perturbation.

**Figure 7.**
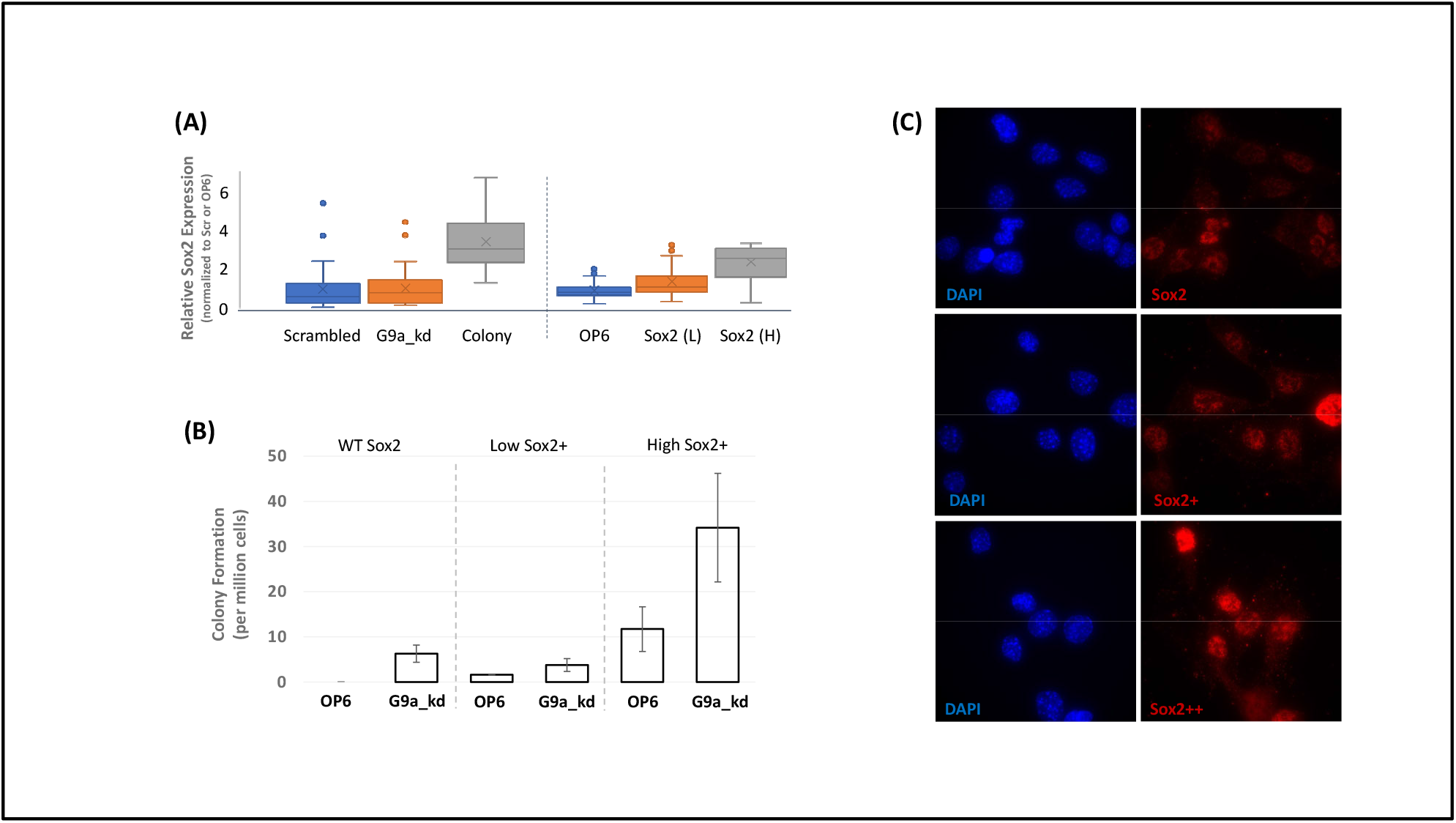
Sox2 expression enables colony formation in OP6 populations. (A) *Left:* Distribution of relative Sox2 antibody intensities Scrambled cells, G9a_kd cells in GFP+ cells from primary transfected OP6 populations. *Right*: Distribution of relative Sox2 expression in OP6 cells and two Sox2 transfections, one with lower overall Sox2 expression (L) and one with higher overall Sox2 expression (H). **(B)** Representative images showing Sox2 expression in untreated OP6 cells (top), the low Sox2 over-expression experiment (middle), and the high Sox2 over-expression experiment (bottom). **(C)** Incidence of colony formation in untreated OP6 cells versus G9a_kd populations (left panel), in OP6 cells with Sox2 over-expression at lower levels, both with and without G9a_kd (middle panel), and in OP6 cells with Sox2 over-expression at higher levels, both with and without G9a_kd (right panel).

## 3 Discussion

The generation of 3-dimensional colonies upon perturbation of G9a protein function in OP6 and OP27 cells was presumably enabled by stochastic Sox2 expression and required progression through the cell cycle. The Sox2 transcription factor protein plays a critical role in somatic cell reprogramming by regulating gene networks that drive differentiated cells back to a pluripotent state (Sommer et al., 2009; Takahashi and Yamanaka, 2006), as well as for retaining pluripotency in neural stem cells (Ellis et al., 2004; Tucker et al., 2010). In the current study, Sox2 expression might have been critical for the up-regulation of either signaling molecules (e.g., TGB1) and/or colony-forming genes, including stem markers.

Although OP6 cells have mostly lost Sox2 expression at the global population level, as have immature neurons of the olfactory lineage (Krolewski et al., 2013), stochastic outliers with elevated Sox2 expression might constitute a bet-hedging strategy in a small subpopulation of differentiating cells in order to enable reallocation of cellular resources within a regenerative regime. Dedifferentiation of neuronal precursor cells within the olfactory epithelium has previously been observed, and strikingly, this dedifferentiation process was compromised in a Sox2 depleted background (Lin et al., 2017). If OP6 colony formation involved Sox2-mediated dedifferentiation, it seems likely that G9a perturbation had shifted the epigenetic landscape to provide greater access to Sox2 target genes, and this shift would presumably depend on transition through the cell cycle in order for these targeted loci to lose H3K9 methylation marks (Ma et al., 2015; Wang and Higgins, 2013). Future work will focus on the role of Sox2 and/or chromatin changes induced within a regenerative context, permit differentiating cells of the olfactory epithelium to reverse their developmental trajectory in vivo.

### 3.1 Colony Cell Diversity in Consideration of Ancestral Models

Since colonies exhibited heterogeneous expression of G9a, consisting of numerous G9a knockdown cells and G9a+ cells, and since we did not observe colony formation after sorting G9a_kd cells from transfected populations, it seems most likely that the transformed OP6 cell colonies derived biclonally through a partnership between a wildtype (G9a+) and G9a_kd cell. Alternatively, it is possible that colonies derived monoclonally from a single founder cell, however, such a model is less parsimonious, requiring that either the colony founder was G9a+ with selection favoring loss of G9a expression during colony growth, or the founder was G9a- and somehow regained robust expression of G9a (e.g., through gene conversion events). Regardless of the underlying origin, the existence of both G9a+ and G9a- cells within colonies suggests that colony development favored G9a heterogeneity. Such a mixed milieu might have been advantageous if colony development depended on some of the cells expressing G9a to silence genes *de novo*, whereas partner cells might have gained some critical functionality as a consequence of G9a knockdown.

We propose a model for colony formation that would seem to be most parsimonious and consistent with all our data (Fig. 8). This model specifies a collaboration between a G9a_kd cell from the majority C1 subpopulation whose gene expression status most resembled the C6 subpopulation of control OP6 cells, partnering with an untransfected OP6 cell going through the cell cycle (i.e., C7-like). These two cells formed a biclonal colony consisting of C3 (G9a-) and C5 (G9a+) cells. This model thus accounts for the mixture of G9a protein expression levels within colonies, as well as the strongly correlated ancestral relationships (C3 with C6/C1 and C5 with C7). Next, the development of a stem-like state within colonies might have been caused by up-regulation of signaling molecules enabled by G9a knockdown within C3 cells (e.g., TGFB1), which might have driven C5 colony cells to transition towards ESC-like states represented by the C4 colony subpopulation. This latter transition would account for PCC similarity between C4 and C5, RNA velocity data suggesting C5 to C4 transitions, and the fact that C4 had maximally diverged from ancestral states and towards ESC-like states. There are likely several models for colony ancestry and progression that would be consistent with all or most of our data, and future work will focus on testing predictions of the partnership model proposed in Figure 8, as well as alternative models. A deeper understanding of this colony formation process will be important for identifying conditions that would favor similar transformation events in vivo.

**Figure 8.**
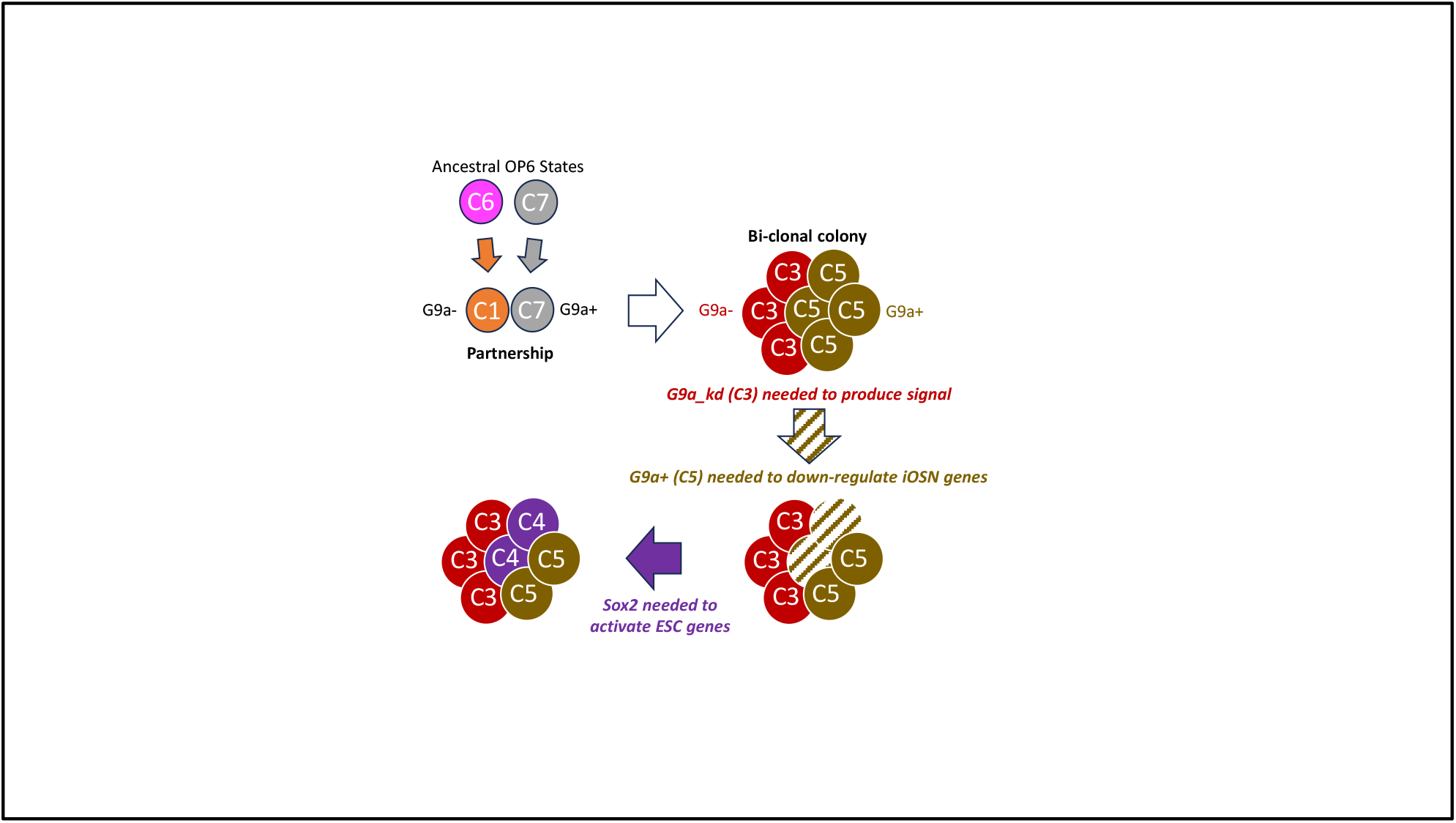
**Model to account for colony cell heterogeneity**. A collaboration between a G9a_kd cell (C6/C1) and a WT OP6 cell (C7/C8) to form a bi-clonal colony with mixed G9a/GFP expression (C3 and C5, respectively), with most divergent C4 colony subpopulation evolving after colony formation via C3 signaling that causes C5 de-differentiation.

### 3.2 Are some of the colony cells de-differentiating to embryonic stem cells (ESCs)?

Regardless of the ancestral histories of various colony subpopulations, we are left with the unanswered question as to whether some of the cells within colonies had been reprogrammed to become pluripotent stem cells. We have described provocative indicators, including growth characteristics, cell morphological changes, chromatin re-organization, iOSN gene down- regulation, and stem-cell marker up-regulation. There are alternative interpretations not yet dismissed, such as the formation of a rogue population of cells that misregulated developmentally associated gene sets in a disorganized way and without proper regulation of the associated gene networks. For example, certain types of tumor cells exhibit global gene misregulation (Fabregat, 2009; Lee and Young, 2013; Zhang and Manley, 2013), including misregulation of stem-cell genes (Nairuz et al., 2023; Zeng et al., 2023). While we have not conducted a full set of experiments to more rigorously investigate tumor properties of OP6 colonies, we do note that these colonies exhibited sensitivity to surrounding cell crowding, thus would seem to retain attributes of contact inhibition (McClatchey and Yap, 2012), and had not attained mitotic independence nor more extreme enhancement of doubling times; these observations suggest more organized as opposed to disorganized growth characteristics. Future studies will focus on further investigation of stem cell-like properties of these OP6-derived colonies, with focus on their pluripotent capacity; e.g., by utilizing strategies commonly employed for validating iPSCs both in vitro and in vivo (Hentze et al., 2009; Nelakanti et al., 2015; Oh and Jang, 2019). We are further inspired by a prior study demonstrating the reprogramming potential of even post-mitotic neurons, a process that required similar criteria involving chromatin perturbation (REST over-expression in this case) alongside activation of stem regulatory networks (Kim et al., 2011). Should transformed OP6 cells meet established criteria in the field for iPSC status, we would further focus investigation as to whether Sox2 and/or G9a manipulations of cells isolated from the olfactory epithelium phenocopy transformed OP6 cells, and thus pursue the possibility that the regenerative environment within the olfactory system could provide a new source of stem cells with neuronal origins.

## 4 METHODS

### 4.1 Cell culturing

The OP6 and OP27 cell lines were used in this study (Illing et al., 2002). We have utilized these lines for decades in studies of olfactory sensory neuronal development. Both lines were grown at 33^0^C with *Dulbecco’s Modified Eagle Media* (DMEM, Gibco) supplemented with *Fetal Bovine Serum* (FBS, Gibco) to a final concentration of 10% by volume, and with *PenStrep* (Gibco) to a final concentration of 1% by volume. The GD25 cell line (Wennerberg et al., 1996) was cultured at 37^0^C in DMEM supplemented with L-glutamine (Gibco), 10% FBS (Gibco), and 1% penicillin/streptomycin (Gibco). The 293T cell line (DuBridge et al., 1987) was cultured at 37^0^C in DMEM supplemented with L-glutamine (Gibco), 10% FBS (Gibco), and 1% penicillin/streptomycin (Gibco). The CCE mouse embryonic stem cell (ESC) line (Robertson et al., 1986) was cultured in DMEM supplemented with L-glutamine (Gibco), 15% FBS (Gibco), 1% penicillin/streptomycin (Gibco), 50uM beta-mercaptoethanol (Sigma) and 1,000 U/mL leukemia inhibitory factor (Invitrogen). In some experiments, we grew cells in media supplemented with 1 mM or 2 mM of the drug BIX01294 G9a inhibitor (Sigma). For passaging, confluent cell cultures were washed twice with PBS before treatment with 0.05% of trypsin (Gibco). Both primary and secondary colonies were transferred by excising colonies with a cell scraper and collecting cells and colonies in 2 mL of media. This media containing colonies was transferred to a 50 mL conical tube, an additional 10.5 mL of media was added, and clumps were vigorously agitated 8-12 times with a 10 mL serological pipet in order to partially disrupt colonies prior transferring to a new 6-well plate. For control cell populations lacking colonies, a random area of the well was scraped off and treated in the same manner as described for colonies.

### 4.2 Western Blot

OP6 cells were seeded in T75 flasks at 20% confluency and treated with either 2 or 4 µM of BIX compound in OP6 media or with OP6 media alone for a duration of 4 days, with media replenished every other day. Following treatment, cells were trypsinized, washed twice with phosphate-buffered saline (PBS), and pelleted. Cell pellets were resuspended in 300- 500 µl of Radio-Immunoprecipitation Assay (RIPA) lysis buffer supplemented with 30-50 µl of 10x protease inhibitor cocktail. The resuspended cells were agitated for 30 minutes on a shaker placed on ice and subsequently centrifuged for 20 minutes at 12,000 rpm. The resulting supernatant containing protein lysates was carefully transferred to a fresh tube. Protein concentrations in the samples were determined using the Pierce BCA Protein Assay Kit following the manufacturer’s instructions. Samples were then denatured by boiling for 5 minutes in 1x Laemmli buffer. Western blotting was performed by electrophoretically transferring the denatured protein samples from a sodium dodecyl sulfate-polyacrylamide gel electrophoresis (SDS-PAGE) gel onto a polyvinylidene difluoride (PVDF) membrane. Post-transfer, the membrane was briefly washed with 1x Tris-buffered saline containing 0.1% Tween-20 (TBST) and then incubated for 30 minutes at room temperature with agitation in 1x TBST containing 5% bovine serum albumin (BSA) to block nonspecific binding. The membrane was subsequently incubated overnight at 4°C with agitation in primary antibody diluted in 1x TBST/5% BSA. Following primary antibody incubation, the membrane was washed three times with 1x TBST and then incubated in secondary antibody diluted in 1x TBST/5% BSA for 1 hour at room temperature. After further washing with 1x TBST, the membrane was developed using NBT/BCIP in alkaline phosphatase buffer. Western blot relative band intensities were quantified using ImageJ (Fiji). Primary antibodies used for Western blotting were as follows: rabbit anti-H3 (Abcam, catalog number ab1791) diluted at 1:5000, mouse anti-H3K9me2 (Abcam, catalog number ab1220) diluted at 1:2000, rabbit anti-H3K9me3 (Millipore, catalog number 07-442) diluted at 1:2000, and rabbit anti-Actin (Abcam, catalog number: Ab8227) diluted at 1:5000. Secondary antibodies used for Western blotting were as follows: donkey anti-rabbit alkaline phosphatase (Abcam, catalog number ab97061) diluted at 1:10,000, mouse anti-rabbit alkaline phosphatase (Sigma, catalog number A2556) diluted at 1:1000, and goat anti-mouse alkaline phosphatase (Sigma, catalog number A3562) diluted at 1:1000.

### 4.3 CRISPR-Cas9 gene targeting

We adopted the CRISPR-Cas9 protocol according to manufacturer protocols (Addgene). Briefly, Cas9 target sites were selected using the sgRNA design tool available at the Broad Institute (http://portals.broadinstitute.org/gpp/public/analysis-tools/sgrna-design). We designed sgRNAs targeting Exon 8 of the G9a gene, as well as a scrambled sequence negative control that should not target endogenous mouse loci (E8 Forward: CACCGATGGCGGAAAGACAGCCCGT, E8 Reverse: AAACACGGGCTGTCTTTCCGCCATC, Scrambled Forward: CACCGGCGAGGTATTCGGCTCCGCG, Scrambled Reverse: AAACCGCGGAGCCGAATACCTCGCC). Complementary oligonucleotides 25 bp in length were synthesized, annealed, and cloned into the lentiviral CRISPR-Cas9 delivery vector containing spCas9-eGFP under control of the EFS promoter and the sgRNA under control of the U6 promoter (Addgene). Following transfection into competent Stbl3 bacteria (Invitrogen), positive transformants were verified by sequencing and plasmids prepared using the Qiagen miniprep kit (Qiagen). Resulting CRISPR/Cas9 lentiviral constructs were transiently transfected along with packaging plasmids pCMV-dR8.2dvpr and pCMV-VSV-G (Addgene) into HEK- 293T cells (ATCC) using standard Lipofectamine LTX transfection protocol (Invitrogen). Virus-containing supernatants were collected 36 and 60 hours post-transfection. Approximately 10^5^ host OP6 cells were subjected to infection at RT in the presence of 8 ug/ml polybrene as described elsewhere (Stewart et al., 2003). Lentivirus-infected OP6 cells/colonies were selected using GFP for subsequent secondary colony analyses.

We measured colony incidence in primary transfected (or BIX-treated) cultures 7-10 days post-treatment. For some experiments, we grew cultures at the non-permissive temperature for the Large T-antigen (39^0^C). Colony growth rates in living colonies were estimated using a standard curve that related maximum cross-sectional colony area to cell numbers, which we derived from a confocal analysis of 16 sample colonies of various sizes. Growth rates (*gr*) of colonies under various experimental conditions were calculated using:

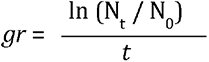

where N_t_ is the number of cells at time *t*, N_0_ is the initial number of cells, and *t* is the elapsed time (Singer, 1986).

### 4.4 Immunocytochemistry and Alkaline Phosphatase staining

Cells and colonies were fixed in 3% paraformaldehyde for 10 minutes and 8% paraformaldehyde for 30 minutes, in PBS buffer containing 0.5% Triton-X (Sigma). Fixed cells were blocked in PBS containing 1% BSA and 0.1% Tween-20 for 20 minutes at 37°C. Primary and secondary antibody incubations were performed at 37°C for 45 minutes in a humidified chamber (except for primary incubations on colonies, which were performed overnight at 4°C). Non-specific antibody interactions were washed in PBS containing 0.1% Tween-20 at 5-minute intervals and with gentle agitation. The following primary antibodies and dilutions were used in this study: rabbit anti-G9a/EHMT2 (C6H3) (Cell Signaling, 3306T, 1:200), anti-histone H3K9me3 (Abcam, ab8898, 1:200), anti-histone H3K9me2 (Abcam, ab1220, 1:500), anti-H4 acetylation (Millipore, 06-866, 1:500), anti- Sox2 (Abcam, ab92494, 1:200), anti-Oct4 (Abcam, ab184665, 1:200), anti-Klf4 (Abcam, ab106629, 1:200), anti-Nanog (Cell Signaling, 8822S, 1:200), anti-cMyc (Cell Signaling, 5605S, 1:200). The following secondary antibodies and dilutions were used: donkey anti-mouse-Cy3 (Jackson Immunoresearch, 715-165-150, 1:100), goat anti-mouse Alexa 488 (Abcam, ab150117, 1:200), donkey anti-rabbit Alexa 488 (Jackson Immunoresearch, 711-545-152, 1:100), and goat anti-rabbit-Cy3 (Millipore, AP132C, 1:800). Cell nuclei were stained with DAPI (1 mg/ml) supplied within the Vectashield mounting solution (Vector Labs). Immunofluorescence images were acquired using a Deltavision RT imaging system (Applied Precision) adapted to an Olympus (IX71) microscope equipped with XYZ motorized stage. Images were processed using Softworx (Applied Precision) or FIJI (ImageJ). Individual nuclei were optically sectioned at 0.5 mm intervals to resolve spatial signal patterns, and in some cases, these confocal images were projected onto a single plane for visualization, following which background signal was subtracted from the reading before quantification of the immunofluorescence signal. Alkaline phosphatase staining was conducted on primary OP6 cell cultures on Day-5 post-transfection using NBT/BCIP stock solution (Roche/Sigma 11681451001) or BCIP/NBT-Blue Liquid Substrate Systems for Membranes (Sigma B3804) in accordance with manufacturer’s protocols.

### 4.5 Sox2 Over-Expression

Sox2 plasmid (pCAG-HA-Sox2-IP), obtained from Shinya Yamanaka (Addgene plasmid # 13459; http://n2t.net/addgene:13459; RRID:Addgene_13459) (Maruyama et al., 2005), was utilized to induce over-expression of Sox2 in OP6 cells. Transfection of the plasmid into OP6 cells was accomplished using Lipofectamine LTX reagent following standard protocols. Puromycin selection (4μM) was employed to establish stable over- expression, with untransfected cells serving as controls until their death confirmed a successful elimination timeline. Immunofluorescence analysis, with primary antibody against Sox2 (Abcam, ab92494, 1:200 dilution) and donkey anti-mouse-Cy3 secondary antibody (Jackson Immunoresearch, 715-165-150, 1:100), was performed on two separate transfections to confirm Sox2 expression, as described previously. Quantitative analysis was conducted using ImageJ software.

### 4.6 Bulk RNA Sequencing and Single nucleus RNA sequencing

Untransfected OP6 cells, along with GFP+ sorted cells with G9a knockout (G9a_kd) and GFP+ sorted scrambled control cells, were cultured until reaching approximately 70% confluency. RNA from young colonies was harvested by directly isolating primary colonies approximately 10 days post-transfection, while RNA from mature colonies was obtained from secondary colonies allowed to grow for approximately 3 weeks post-transfection. Total RNA was then extracted from these cells using the Direct-zol RNA Microprep Kit (Zymo Research), following the manufacturer’s instructions. RNA quality and quantity were assessed, and high-quality samples were sent to Novogene for library preparation and RNA sequencing (RNA-seq) analysis according to standard protocols. For single nucleus RNA sequencing, GFP+ sorted cells with G9a knockout (KO) and GFP+ sorted scrambled control cells, along with young colonies harvested by directly isolating primary colonies approximately 10 days post-transfection, were frozen overnight in a medium containing 90% fetal bovine serum (FBS) and 10% dimethyl sulfoxide (DMSO) and sent to Novogene for single nucleus RNA sequencing and subsequent analysis. GO analysis was conducted by Novogene, providing the top ten GO terms from each category (Biological Process, Cellular Component, and Molecular Function).

### 4.7 Bioinformatics analyses of RNA-seq data

Raw fastq files were processed using 10X Genomic’s Cellranger pipeline (v 7.2.0), mapped using the standard configuration onto a modified GRCm39 genomic annotation build that had GFP and WPRE elements added as distinct transcripts. Mapped BAM files were then count processed using velocyto v0.17.8 (La Manno et al., 2018) with rmasks enabled to output loom files. Scanpy v 1.9.4 (Wolf et al., 2018) was used for downstream processing. Data was filtered on a minimum of 200 genes per cell, 2 cells per gene, and 600 genes by counts. Predicted doublets were removed using Scrublet (Wolock et al., 2019). Integration of the three datasets was done using the bbknn algorithm built into Scanpy. RNA velocity was estimated using scvelo (La Manno et al., 2018) and the deterministic algorithm. Pseudobulk counts of each cluster was generated using the get_pseudobulk algorithm found in the decoupler package (Badia et al., 2022). Pseudobulking of single-cell RNA sequencing (scRNA-seq) data was performed to aggregate unique molecular identifier (UMI) counts within a specific cell cluster. This process involved summing UMI counts for each gene across individual cells within the cluster, followed by normalization of the total counts to 1 million. Pseudobulking facilitated comprehensive analysis of gene expression profiles within distinct cellular populations, providing insights into their biological characteristics and functional roles.

In some analyses, we compared pseudobulked single-cell sequenced subpopulations to ESC (Kinoshita et al., 2021) and iOSN (Zunitch et al., 2023) transcriptomes by first identifying a subset of genes that were at least 10-fold up- (UP set) or down-regulated (DOWN-set) in ESC versus iOSN datasets (and vice versa), with a minimum expression level of 1.0 FPKM in the cell type with higher expression of the two. We then calculated the average FPKM expression level in a sequenced sample for the UP and DOWN gene set. A net ESC FPKM score was derived by subtracting the average FPKM for the DOWN ESC set from the UP ESC set (similarly, a net iOSN FPKM score was derived by subtracting the average FPKM for the DOWN iOSN set from the UP iOSN set).

For some comparative analyses, we derived a weighted scoring system to assess signature gene profiles within sequenced samples. For these analyses, we used ranked signature gene lists for a given cell type (Zunitch et al., 2023). Signature genes were obtained through the FindMarkers algorithm implemented in Scanpy on the cell populations identified in (Zunitch et al., 2023). We assigned a linearized weight between 0-1 according to rank of each list. A total score for a sequenced sample against a particular signature gene list was calculated based on the sum of individual weighted FPKM levels as follows: S (*FPKM Score*) = S (Normalized FPKM Level * Relevance Weight).

## ACKNOWLEDGMENTS

This work was supported by National Institutes of Health Grant R01- DC006267. The authors wish to thank Jane Roskams for generously providing the OP6 and OP27 cell lines. We extend our appreciation to James Schwob and Chris McGinnis for insightful discussions.

## 5. COMPETING INTERESTS

No competing interests declared.

## Notes

### Competing Interest Statement

The authors have declared no competing interest.

